# Population and community variability deviate from stationary expectations during transient dynamics

**DOI:** 10.64898/2026.07.08.737188

**Authors:** Josquin Guerber, Damien Genettais, Colin Fontaine, Elisa Thébault

## Abstract

Under complex perturbation regimes, biodiversity dynamics show temporal variability in species and community abundance around long-term population trends. Many species indeed show long-term declines while other species increase, putting natural communities far from stationary regimes, while variability is often studied near equilibrium. We contribute to bridging this gap by investigating population and community variability during long-term trends caused by press perturbations in stochastic models of population dynamics. By estimating the deterministic changes in mean and variance during the transient regime, we show that population variability deviates from stationary expectations. Moreover, the deviation strongly depends on the sign of the population trends: increases generate excesses of variability while declines generate deficits. Scaling up to community variability, we propose a decomposition of community variability deviation, allowing to highlight that community variability in the transient regime depends on how the press perturbation is distributed within species relative abundances and growth rates. These results challenge the equilibrium assumption and open new perspectives for the study of the variability of ecological systems under multiple perturbation types.

## Introduction

Understanding how ecological systems respond to perturbations is critical in assessing threats to the maintenance of ecosystem functioning (Cardinale et al. 2012). Shortterm fluctuations in environmental conditions generate variable population and community abundances (Tilman 1996; Engen et al. 1998), and anthropogenic pressures cause many wild species to decline in abundance while other species increase (McKinney and Lockwood 1999; Dornelas et al. 2014). While empirical and experimental approaches have proposed different ways to disentangle temporal variability from so-called long-term population trends (Hillebrand et al. 2018; Lepš et al. 2019; Clark et al. 2022), theoretical understanding of how temporal variability might be affected by long-term population trends is lacking. In some situations, these two components of ecological stability appear correlated (Donohue et al. 2013), but no underlying mechanisms for such correlation have been proposed.

Temporal variability encapsulates both the intensity of environmental stochasticity as well as the ecological system’s intrinsic ability to recover from perturbations (Arnoldi et al. 2019; Van Meerbeek et al. 2021). Because of this dual property as well as its ease of measurement, temporal variability has received considerable theoretical and empirical attention (Kéfi et al. 2019). There, temporal variability is often measured as the coefficient of variation, the ratio of temporal standard deviation to temporal mean, yielding an adimensional index of how variable a system is compared to its magnitude (Tilman 1996). A common framework for investigating temporal variability is to think of small perturbations affecting a system close to its equilibrium (Lundberg et al. 2000; Arnoldi et al. 2019). Following this near-equilibrium assumption, most studies rule out the impact of long-term population trends on the measured variability by studying a stationary regime (Arnoldi et al. 2019; Danet et al. 2025) or by detrending time series prior to variability analysis (Craven et al. 2018; Li et al. 2021). Under anthropogenic pressures, long-term population trends challenge the near-equilibrium assumption, with shifts in community composition being the most empirically supported aspect of recent biodiversity changes at local scales (Carroll et al. 2023; Dornelas et al. 2023). While the importance of non-equilibrium dynamics is increasingly recognized in ecological theory (Hastings et al. 2018), they are not often combined with the study of temporal variability. A reason for this theoretical gap lies in the fact that the most commonly used mathematical methods for studying the stability of dynamical systems require to linearize the system’s dynamics around the equilibrium, which is only reasonable if the system stays in the close vicinity of its equilibrium.

The near-equilibrium assumption has yielded a general theory of how temporal variability scales across spatio-temporal and ecological dimensions (Clark et al. 2021). Temporal variability of population and community abundance has been shown to vary with the strength of stochasticity (Arnoldi et al. 2019) and with species growth rate (Mentges et al. 2024). Moreover, the near-equilibrium assumption has also enabled the development and mechanical understanding of the insurance hypothesis (Yachi and Loreau 1999), which states that the variability of community abundance or biomass declines with the number of species in the community (Tilman 1996; Pennekamp et al. 2018). Community abundance is more variable when the species’ population abundances are more variable, or when the species-level fluctuations are synchronized and fail to compensate for each other (Loreau and de Mazancourt 2008; Thibaut and Connolly 2013). Biodiversity has been shown to promote the evenness of species variances, decreasing species synchrony and thus stabilizing communities (Zhao et al. 2022), while interspecific competition tends to increase population variability (Ives et al. 1999). Understanding how these mechanisms could operate far from equilibrium is a critical step towards better understanding the temporal dynamics of natural communities.

In this study, we implement long-term trends in stochastic population and community dynamics models as responses to press perturbations, *i*.*e*. sudden and maintained changes in carrying capacity (Van Meerbeek et al. 2021; Lajaaiti et al. 2025). Our approach relies on deterministic approximations for the changes in the mean and variance of a process described by a stochastic differential equation, called moment closure approximations (Särkkä and Solin 2019). In the biochemical literature, these approximation methods have enabled exploration of the role of stochastic phenomena in cell biology (Schnoerr et al. 2017), including in the case of transient dynamics (Zechner et al. 2012). While they have been known for some time in ecology (Lande et al. 2003; Nåsell 2003), moment closure approximations are rarely used in the context of population dynamics models. Here we use such approximations in order to summarize stochastic dynamics during the long-term trends into simple, deterministic indices for population and community variability far from equilibrium.

We revisit classical findings on temporal variability of population and community abundance in the context of long-term trends caused by press perturbations. At the population level, we investigate how properties of the press perturbation and population growth rate drive temporal variability during long-term trends. We then explore how the variability of a simple community is affected by the diversity of individual species’ responses to the press perturbation and by how these responses depend on species abundances and growth rates, this with and without interspecific competition. This work extends current knowledge on population and community variability to contexts with transient dynamics and paves the way for better inference of ecological stability and perturbation regime from variability data.

## Methods

### Mathematical framework

In a community of *S* species under environmental stochasticity (Engen et al. 1998; Lande et al. 2003; Arnoldi et al. 2019), population abundances are the result of community dynamics (drift) and stochastic pulse perturbations (diffusion). In continuous time, this translates to some stochastic differential equation system of the form:

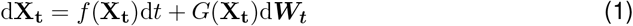

**X**_**t**_ = (*X*_1_(*t*), *X*_2_(*t*), …, *X*_*S*_(*t*)) is the vector of population abundances at time *t, f* encapsulates the deterministic community dynamics and *G* represents how small short-term perturbations modelled as infinitesimal Wiener process increments d**W**_**t**_ affect the different species. For simplicity, we will call *f* the deterministic term and *G* the stochastic term, but notice that *G* itself is not random and only *G*(**X**_**t**_)d***W***_***t***_ is.

For any stochastic process that can be written as Equation 1, stochastic calculus allows to approximate the changes in the statistical moments of **X**_**t**_ as an ordinary, deterministic differential equation system (see supplemental Section S1 for an overview). In particular, our moment closure approximation relies on linearizing the deterministic term around the (time-varying) mean of the stochastic process instead of linearizing it around a fixed point (Särkkä and Solin 2019). The approximation thus does not only hold in a stationary regime near an equilibrium, but also during transient regimes. Moreover, the mean and variance of species abundance can be studied as numerical solutions of ordinary differential equations instead of the sample mean and variance of many simulations of the stochastic process (Figure 1).

**Figure 1:**
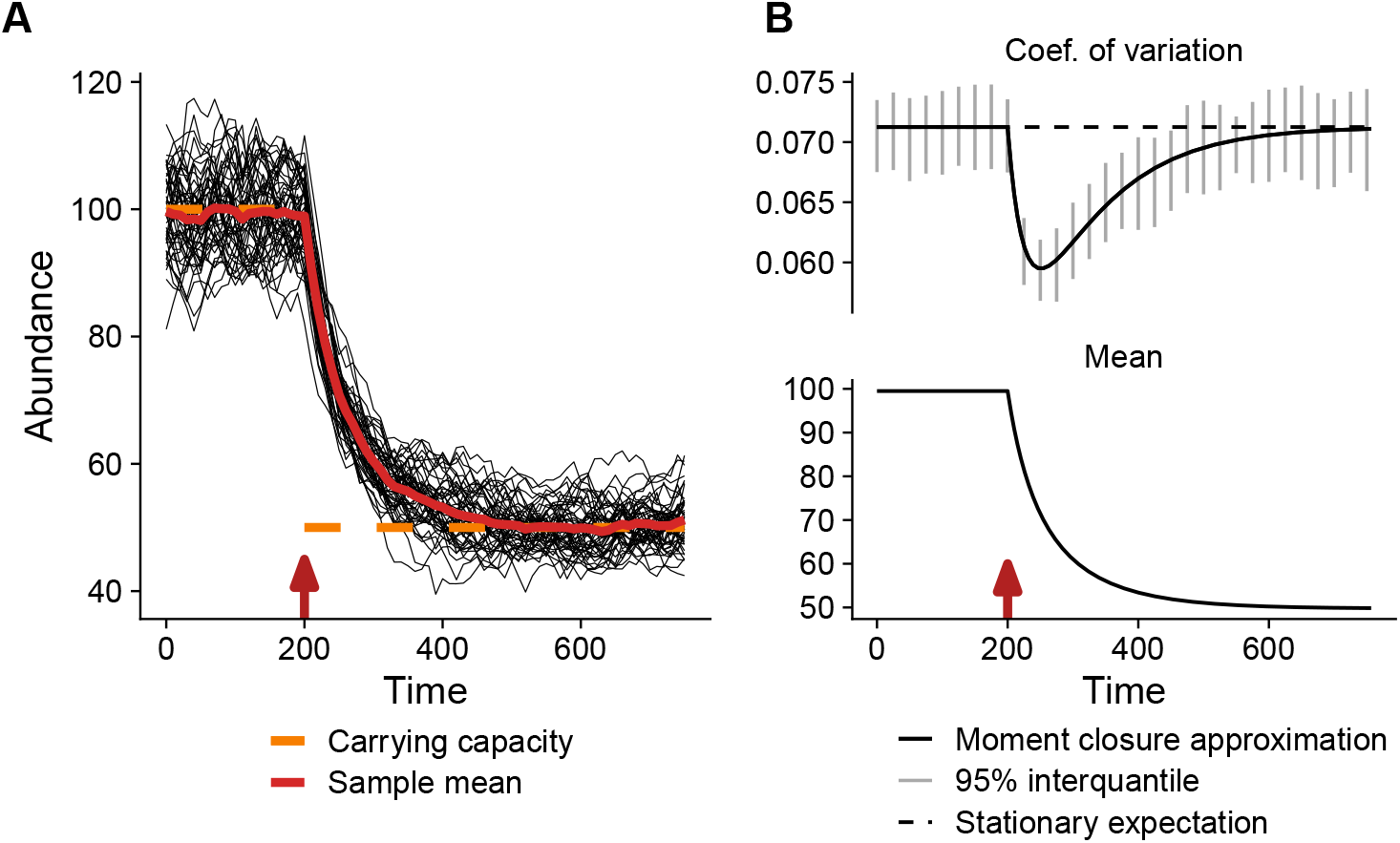
Population dynamics with a stochastic logistic growth under a press perturbation. (A) 50 numerical simulations of the stochastic logistic growth with a press perturbation from *t*_*d*_ = 200 (red arrow) onwards. (B) Solution of the moment closure approximation for coefficient of variation (top) and mean (bottom). Gray vertical intervals on the top panel indicate the 95% interquantile for 50 replicates of the coefficient of variation computed across 500 stochastic simulations. The corresponding intervals for the mean had negligible sizes. The dashed black line indicates the stationary expectation for the coefficient of variation. *K*_0_, *K*_*d*_, *r* and *σ*_*e*_ were set to 100, 50, 0.01 and 0.01, respectively.

### Mean and variance of a single population

For a single population with a carrying capacity *K* and a logistic growth rate *r* under environmental stochasticity of strength *σ*_*e*_, Equation 1 simplifies to:

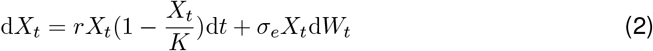

Under the moment closure approximation (supplemental Section S1.2), the dynamics of the mean and variance of population abundance, *µ*_*t*_ and *V*_*t*_, can be described as:

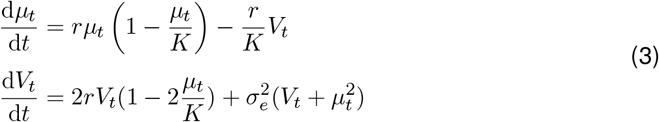

System 3 holds during transient regimes but also allows setting 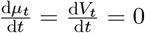 in order to calculate (*µ*^∗^, *V* ^∗^), the mean and variance of *X*_*t*_ in the stationary regime. Conversely, for any observed mean value *µ*_*t*_, the same system can be solved for 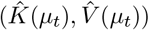, with 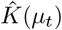 the carrying capacity required to observe a stationary regime of mean 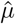, with variance 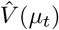. We call 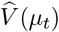 the stationary expectation for variance at mean *µ*_*t*_, it is the variance that the population would have if it was in a stationary regime at mean *µ*_*t*_.

A given population of mean *µ*_*t*_ thus has three associated values for variance: *V*_*t*_, the observed variance according to the moment closure approximation; 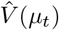, the stationary expectation for variance at *µ*_*t*_; and *V* ^∗^, the variance in the stationary regime. Note that 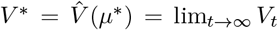. Because *V*_*t*_ (resp. *V* ^∗^) is always roughly proportional to 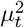 (resp. *µ*^∗2^), we compute variability as the coefficient of variation of *X*_*t*_, *i*.*e*. 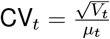 (see supplemental Section S2.1 for derivation).

### Long-term trends as responses to press perturbations

We model long-term population trends as the transient response of the population to a press perturbation, *i*.*e*. to a sudden and maintained change in carrying capacity (Lajaaiti et al. 2025). Press perturbations that decrease the carrying capacity generate decreasing population trends, while press perturbations that increase *K* generate increasing trends.

Starting from a stationary regime with *K* = *K*_0_ for time *t < t*_*d*_ corresponding to mean and variance 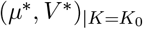, we introduce a press perturbation by setting *K* = *K*_*d*_. The relative strength of the press perturbation is defined as 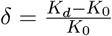. A single simulation of Equation 2 shows how a single replicate population reacts to the press perturbation (one single black line in Fig. 1A). We first assess the approximation provided by Equation 3 for the mean and variance of population abundance as it responds to a press perturbation. To do so, we compare the (deterministic) expected mean and variance provided by Equation 3 with the distribution of observed mean and variance obtained from simulating Equation 2. To estimate these distributions across the transient regime, we first compute the mean and coefficient of variation of a single batch of simulations (e.g. 50 simulations of population dynamics in Figure 1) for each time step. This measures the changes in mean and variability of populations in this batch across time. Second, the distributions of mean and coefficient of variations at each time step can be estimated by simulating many batches, called replicates, and are then compared to their approximate expected values after converting variances from Equation 3 to variabilities (Figure 1B).

Over the course of transient dynamics following the press perturbation, *X*_*t*_ will converge to a new stationary regime 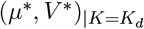 if the population does not become extinct and if the parameter values allow the moment closure approximation. For all values of *µ*_*t*_ during the transient dynamics, we compute a stationary expectation for variability 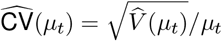, which corresponds to the coefficient of variation that the population would have if it was in a stationary regime at mean *µ*_*t*_. The deviation of the variability of *X*_*t*_ from a stationary regime at a given time can be computed as the log-ratio between CV_*t*_ and 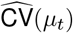. This deviation can then be integrated over the course of the transient regime:

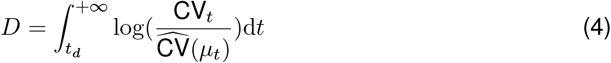

Positive values of *D* indicate an overall excess of variability over the transient regime compared to stationary expectations, while negative values indicate an overall lower variability.

### Community variability in the transient regime

For community dynamics, we set the deterministic term in Equation 1 as the generalized Lotka-Volterra system (Lotka 1925; Volterra 1931). We use its formulation from Bunin (2017):

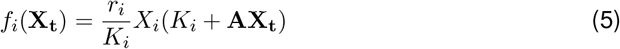

The interaction matrix **A** encapsulates each species’ effect on the abundances of all species (including itself). We set *A*_*ii*_ = −1 for all *i* to ensure that, without any interspecific interactions, each species reaches its carrying capacity *K*_*i*_. Off-diagonal terms can be set at will or randomly chosen, but only some ranges or distributions will allow for asymptotic coexistence (Bunin 2017; Barbier et al. 2018). With environmental stochasticity acting independently on each species with constant strength, the stochastic term is *G*(**X**_**t**_, *t*) = *σ*_*e*_ diag(**X**_**t**_).

We approximate Equation 5 by a deterministic differential equation system for ***µ***_***t***_ and **Σ**_***t***_, the vector of means and the variance-covariance matrix of **X**_**t**_ (supplemental Section S1.2, Equation S5). Because **Σ**_***t***_ is symmetric, this is a system of *S* + *S*(*S* + 1)*/*2 = *S*(*S* + 3)*/*2 equations. Therefore, while the system remains numerically solvable, it quickly becomes analytically untractable as the number of species increases. For any parameter combination, the means and variance-covariance matrix in the stationary regime, (***µ***^**∗**^, **Σ**^**∗**^), can be computed numerically and for any observed value for means ***µ***_***t***_, one can try to compute the carrying capacities 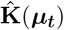 that allow for a stationary regime with populations means ***µ***_***t***_. The corresponding variance-covariance matrix is called 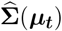, the stationary expectation for the variance-covariance matrix at means ***µ***_***t***_. For some values of ***µ***_***t***_, solutions for 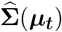 may not exist (in these cases, because of the values of the other parameters, there is no 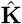 such that a stationary regime is possible at this means).

Community mean *µ*_*c*_(*t*) and variance *V*_*c*_(*t*) can be obtained by summing all elements of ***µ***_***t***_ and **Σ**_***t***_, respectively. Moreover, the stationary expectation for community variance, 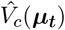, can be computed by summing the elements of 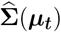. At a given vector of population means 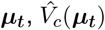 is the variance that the community would have if the system was in a stationary regime at populations means ***µ***_***t***_. The total deviation in community variability from stationary expectation across the transient regime, *D*_*c*_, can thus be defined similarly to Equation 4:

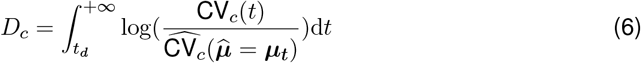

In order to pinpoint the underlying drivers of *D*_*c*_, we use two subsequent decompositions of the community coefficient of variation 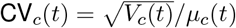. From Thibaut and Connolly (2013), 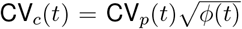, with CV_*p*_ the average of species coefficients of variation weighted by species’ relative abundances and *ϕ* the species synchrony index of Loreau and de Mazancourt (2008). Moreover, following Zhao et al. (2022) and with *s*_*ii*_(*t*) the variance of species *i* (*i*.*e* the *i*-th diagonal element of **Σ**_***t***_),

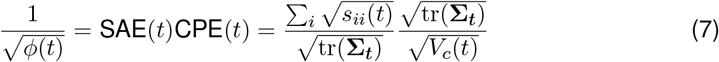

The statistical averaging effect (SAE(*t*)) accounts for how aggregating species with uncorrelated fluctuations reduces overall community variability, and the compensatory dynamics effect (CPE(*t*)) accounts for the reduced variability introduced by negative temporal covariances among species (Zhao et al. 2022). By definition, communities with uneven species variances have a low statistical averaging effect. For each of CV_*p*_(*t*), SAE(*t*) and CPE(*t*), the corresponding expectation under a stationary regime at means ***µ***_***t***_ can be computed from 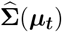. Each component of community variability is then assigned a total deviation across the transient regime, and all their deviations sum to the total deviation in community variability:

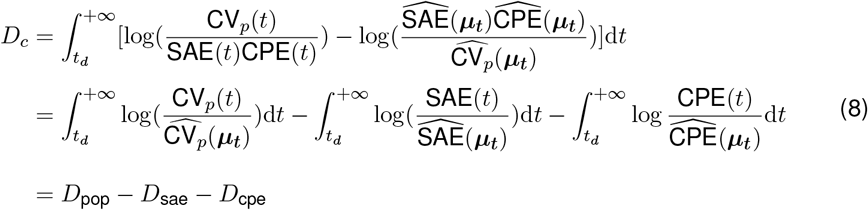

Community variability thus deviates from stationary expectations either because mean population variability, statistical averaging or compensatory dynamics deviate from their stationary expectations. Excesses of statistical averaging effect or of compensatory dynamics effect generate deficits of community variability.

### Press perturbation scenarii and numerical analysis

To explore the deviation of population variability from stationary expectations, we explore the effects of the relative press perturbation *δ* and the population growth rate *r* on *D* by numerically solving equation 3. For all values of *µ*_*t*_ across time in the numerical solution, we compute the stationary expectation for population variability 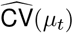 using system 3, which allows to compute *D* (Equation 4) by numerical integration between perturbation time *t*_*d*_ and the end of simulation time.

In order to investigate the impact of press perturbations on community variability, we focus on a community with two species with 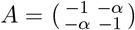, *i*.*e*. symmetric competitive interactions of strength *α*. The vector of carrying capacities before the press perturbation, **K**_**0**_ = (*K*_0,1_, *K*_0,2_) is set such that the *K*_0,1_ *> K*_0,2_. The first species thus is the most abundant in the community before the press perturbation. Together with the vector of growth rates **r**, *α* and **K**_**0**_ define the community context in which the press perturbation is introduced. We used community contexts with or without interspecific competition and with equal or different growth rates across species. Since higher population growth rate is generally associated with higher carrying capacity (Brown et al. 2004), *r*_1_*/r*_2_ was set equal to *K*_0,1_*/K*_0,2_ for contexts where growth rates were different across species.

Setting the relative press perturbation to be equal in magnitude |*δ*| across species, there are four different ways to introduce a press perturbation, which we call perturbation scenarii. First, the two species can have the same response to the press perturbation, which can be negative or positive. This defines the “increase” scenario with ***δ*** = (|*δ*|, |*δ*|) and the “decrease” scenario with ***δ*** = (−|*δ*|, −|*δ*|). In the two other scenarii, the response to the press perturbation varies across species. The “divergence” scenario is defined with ***δ*** = (|*δ*|, −|*δ*|), *i*.*e*. the carrying capacity of the abundant species increases and the other decreases. Conversely, the “convergence” scenario is defined with ***δ*** = (−|*δ*|, |*δ*|), the carrying capacity of the more abundant species is decreased and the carrying capacity of the less abundant species is increased.

We applied the four perturbation scenarii to the community contexts using the moment closure approximation, by numerically solving equation S5. For each combination of perturbation scenario and community parameters, we recorded species means and variance-covariance matrices across time and computed the stationary expectations for the community variability components at each time step, which we then summarized using *D*_*c*_, *D*_pop_, *D*_sae_ and *D*_cpe_ (Equation 6).

Symbols used throughout the Methods section are summarized in Table 1. Stochastic differential equation systems were simulated in the Julia programming language using the `DifferentialEquations.jl` framework (Datseris 2018) and the SRIW1 algorithm (Rößler 2010) with a fixed time step of d*t* = 1*/*25 and *t* from 0 to 1000 with press perturbations at *t*_*d*_ = 200. Ordinary differential equation systems were solved using the `scipy` suite for the Python language, with an adaptive time step. Over all numerical simulations, parameter values were chosen in order to observe noticeable long-term trends after press perturbations while also ensuring (i) that replicate populations did not go extinct for stochastic simulations and (ii) that a stable fixed point of Equation 3 existed before and after the press perturbation for the chosen parameters. Results were visualised using the `ggplot2` package in the R programming language. The manuscript was written as a `Quarto` project.

**Table 1:** List of symbols. Non-bold symbols in parentheses are the one-dimensional counterparts of each bold symbol. They are used for the stochastic logistic growth of single populations while bold symbols are used for the generalized Lotka-Volterra system modelling community dynamics. In the column “Dimension”, for functions, the left-hand side of the arrow is the dimension of the domain of the function and the right-hand side is the dimension of its destination set. Throughout the text, symbols with a ^ hat denote the stationary expectation for their measured quantity.

| **Symbol** | **Description** | **Dimension** |
| --- | --- | --- |
| $S$ | Number of species | scalar |
| $\mathbf{X}_t$ ($X_t$) | Species abundances at time $t$ | $S$ (1) |
| $\boldsymbol{\mu}_t$ ($\mu_t$) | Population means at time $t$ | $S$ (1) |
| $\boldsymbol{\Sigma}_t$ ($V_t$) | Variance-covariance matrix of abundance at time $t$ (variance at time $t$) | $S \times S$ (1) |
| $\mathbf{r}$ ($r$) | Species growth rates | $S$ (1) |
| $\mathbf{K}$ ($K$) | Species carrying capacities | $S$ (1) |
| $\sigma_e$ | Strength of environmental stochasticity | scalar |
| $f$ | Deterministic dynamics term | $S \rightarrow S$ |
| $G$ | Stochastic dynamics term | $S \rightarrow S \times S$ |
| $\mathbf{CV}_t$ | Coefficient of variation of population abundance at time $t$ | scalar |
| $D$ | Total deviation in population variability | scalar |
| $\mathbf{CV}_c(t)$ | Coefficient of variation of community abundance at time $t$ | scalar |
| $D_c$ ($D_{\text{cmp}}$) | Total deviation in community variability (in community variability component cmp) | scalar |
| $t_d$ | Time point where the press perturbation is introduced | scalar |
| $\delta$ ($\delta$) | Relative strength of the press perturbation | $S$ (1) |

## Results

### Population model

During the transient regime after a press perturbation, population variability deviates from its stationary expectation (Figure 1). This is noticeable over many samples from the stochastic logistic growth, and the moment closure approximation allows to reproduce this effect in a deterministic setting (Figure 1B). With the parameters used to produce Figure 1, this deviation is a deficit of variability, and the coefficient of variation reaches down to a value 10% lower than its value if the population was in a stationary regime.

Positive press perturbations, *i*.*e*. press perturbations that increase the carrying capacity, generate excesses of population variability during the positive transient trends in abundance (Figure 2A, green lines). Conversely, negative press perturbations generate deficits of variability during the negative long-term trends (Figure 2A, orange lines). Equation 3 allows to construct the phase space for mean and variance of population abundance after the press perturbation (supplemental Section S2.1). The relative position of the stationary expectation for variance (dashed line on Figure 2B) and of the variance isocline (blue line on Figure 2B) explain that an increase (resp. decrease) in mean comes with a stronger increase (resp. decrease) in variance, ultimately causing an excess (resp. a deficit) of coefficient of variation. Since variability thus does not depend only on the mean abundance but also on the press perturbation, observed variability (solid thin lines on Figure 2B) deviates from its stationary expectation. In other words, during transient dynamics, population variability does not stay at its stationary expectation dictated by the mean abundance, but also depends on the sign and strength of the press perturbation that generated the transient dynamics. This results in a deviation from the stationary expectation for variability. Mathematical analysis justifies that the excess (resp. deficit) of population variance for positive (resp. negative) press perturbations is a general result that does not depend on parameter values (supplemental Section S2.2).

**Figure 2:**
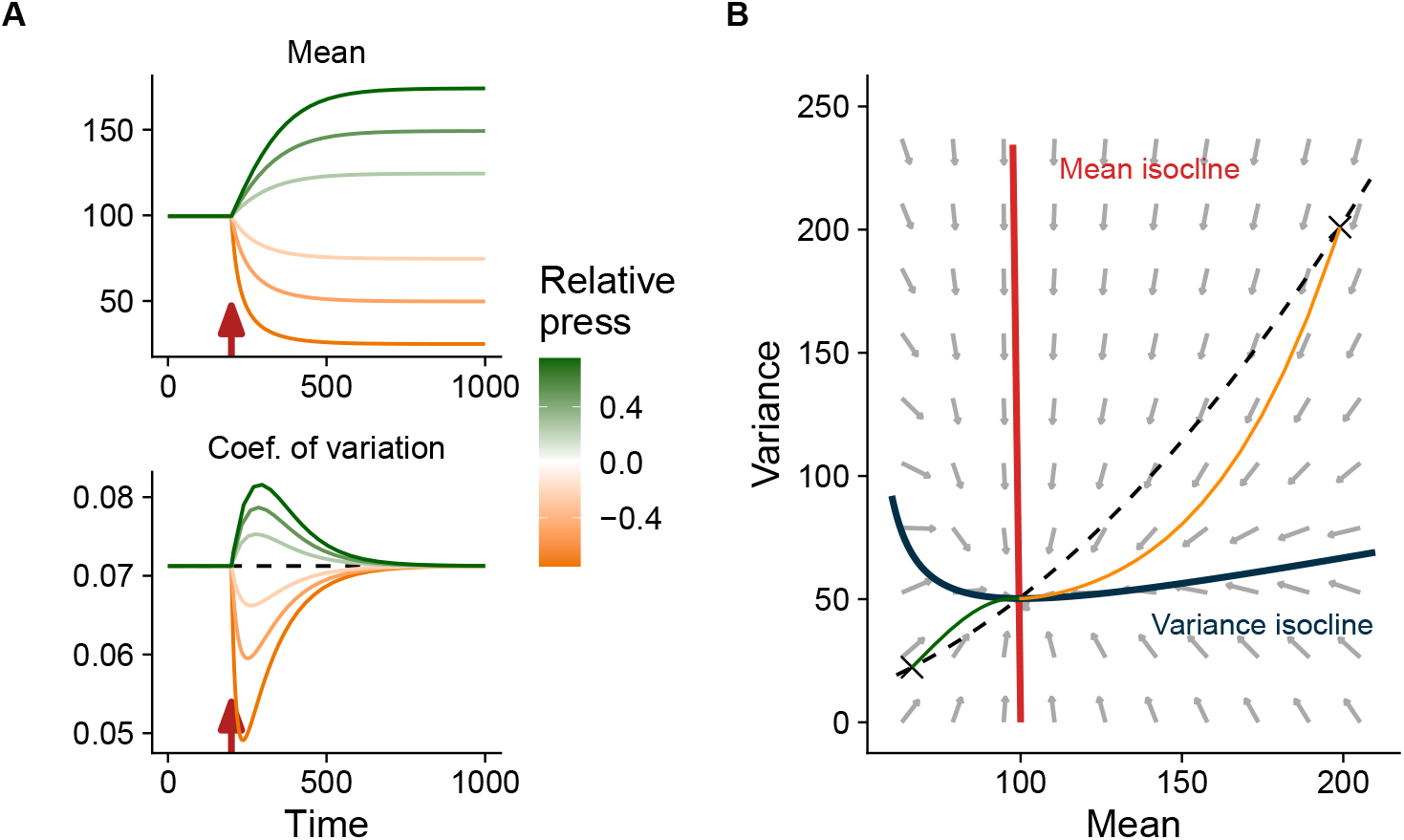
Change in mean and coefficient of variation of population abundance with the sign of the press perturbation for the stochastic logistic growth. (A) In the moment closure approximation, different press perturbations initiated from *K*_0_ = 100 at *t*_*d*_ = 200 (red arrows) induce different responses of mean and coefficient of variation (colored lines). The dashed line shows the stationary expectation for variability. (B) Deterministic phase space for variance and mean for *K*_*d*_ = 50. The dashed line shows the stationary expectations for variance at each possible mean value. The two crosses are the initial conditions for the green and yellow lines (example trajectories respectively for a population increase and decrease). Grey arrows show the direction of the instantaneous dynamics for mean and for variance across the phase space. *r* and *σ*_*e*_ were both set to 0.01.

Integrating the deviation in population variability over the whole duration of the transient regime confirms that the magnitude of the deviation depends on the relative strength of the press perturbation and not on initial carrying capacity (Figure 3A). Comparing absolute values, for a given strength of press perturbation, deviations appear stronger for declines than for increases. Regardless of its sign, total deviation in variability is amplified by stronger environmental stochasticity and buffered by increasing population growth rate (Figure 3B).

**Figure 3:**
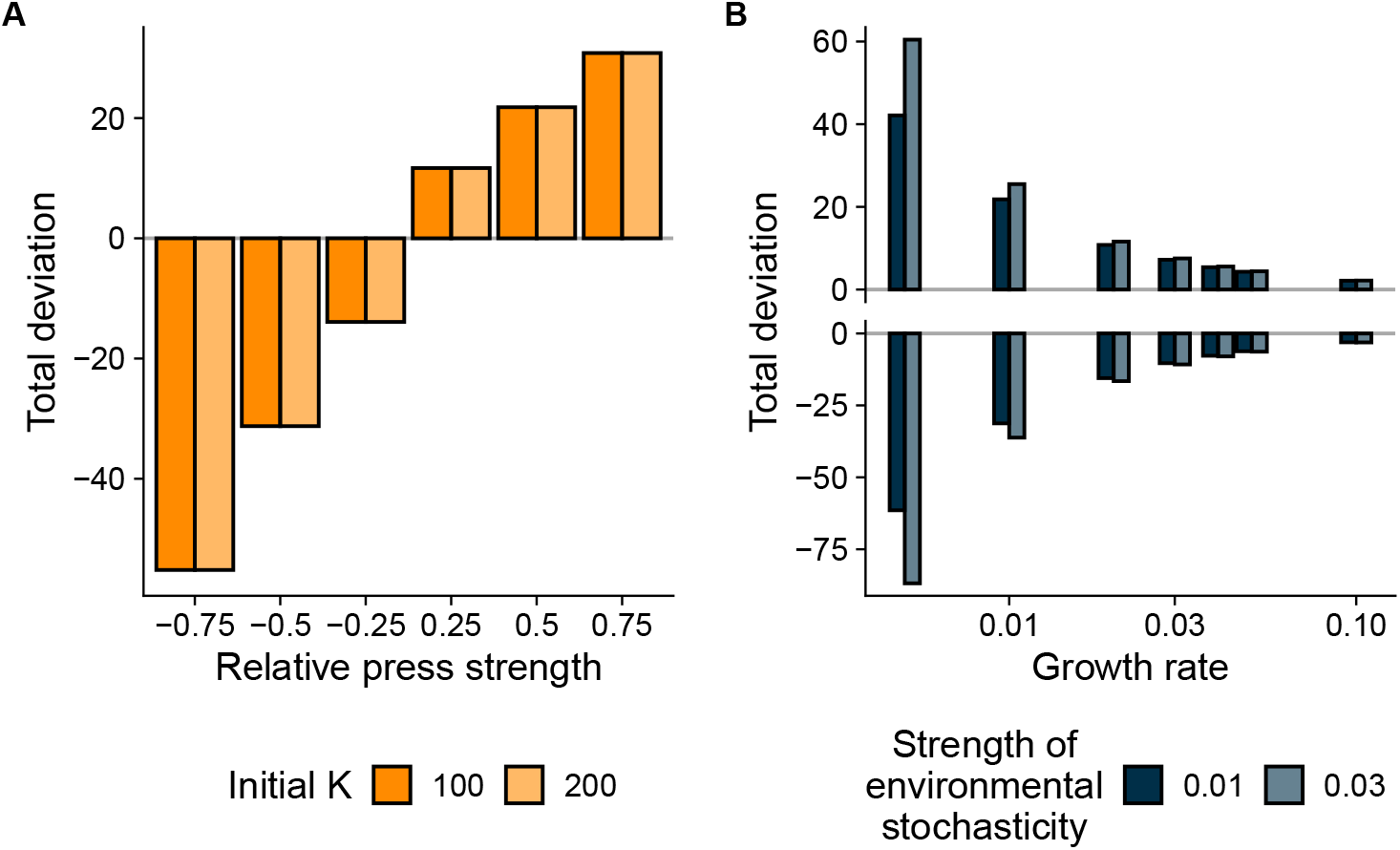
Response of total deviation in population variability from its stationary expectation for the stochastic logistic growth. (A) Total deviation is measured for different relative press strengths *δ* and initial carrying capacities *K*_0_ and (B) for different values of the species growth rate *r* and environmental stochasticity strength *σ*_*e*_. Panel (A) is generated with *r* and *σ*_*e*_ set as 0.01 and panel (B) is generated with *K*_0_ as 100 and *δ* as 0.5 (top part) or -0.5 (bottom part). The horizontal axis of panel (B) is log-scaled.

### Community model

The moment closure approximation for mean and variance combined with the decomposition of deviation in community variability allows to study community variability during the transient regime after a press perturbation (Figure 4). In this example, a community of two species with uneven carrying capacities and moderate interspecific competition undergoes a press perturbation with the “divergence” scenario. At the population level, the rarer species has a strong negative deviation in population variability and the more abundant species has a less intense positive deviation in population variability (Figure 4A). However, the deviation of the most abundant species dominates and community variability during the transient is higher than its stationary expectation.

**Figure 4:**
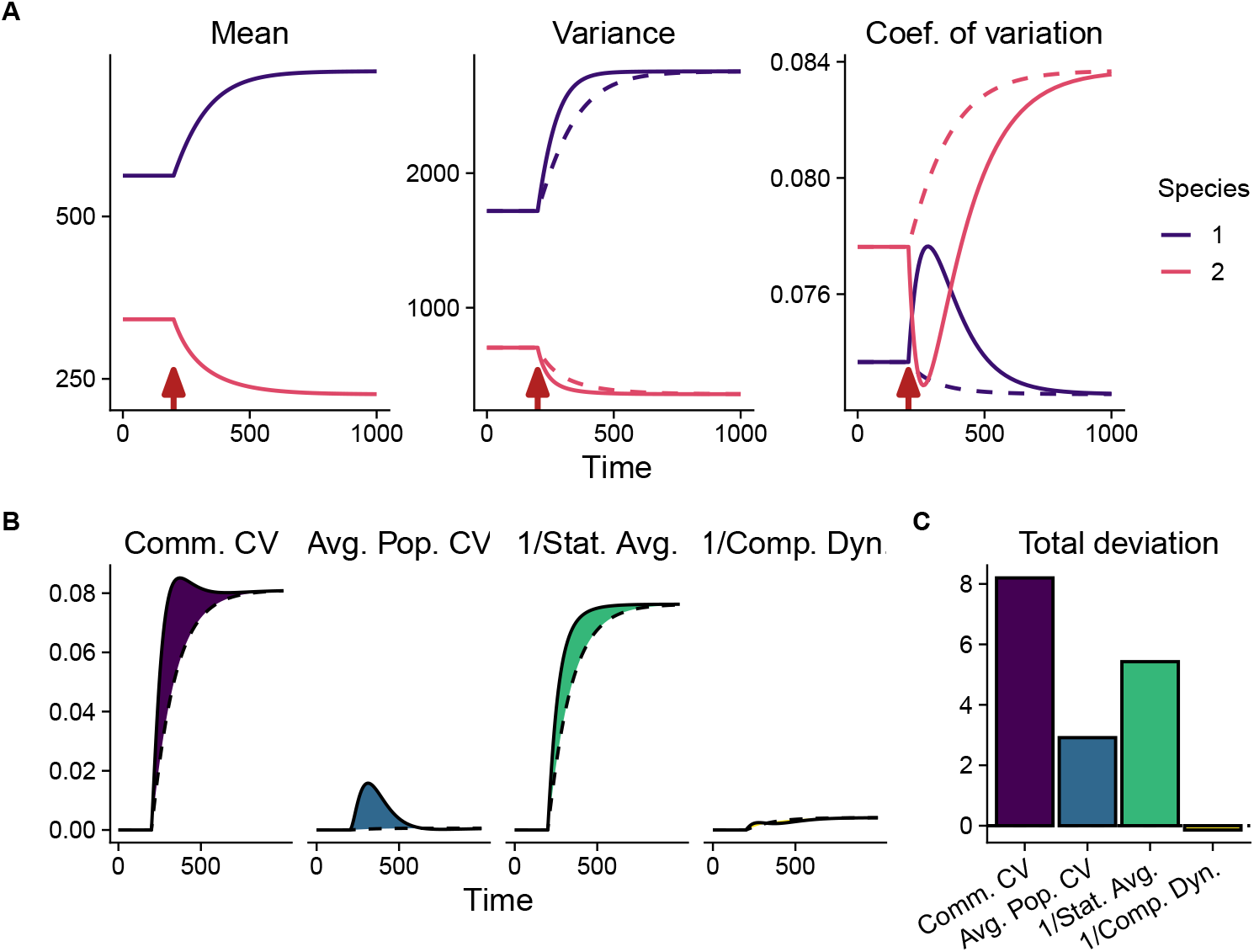
Decomposing community variability during transient dynamics following a press perturbation. (A) Moment closure approximation for means, variances and coefficients of variation for a community of two species undergoing a press perturbation at *t*_*d*_ = 200 (red arrows). Dashed lines indicate, for a given timepoint, the stationary expectations for variances or variabilities (see Methods). (B) Community variability components (solid lines) and their stationary expectations (dashed lines), computed from the moment closure approximation. Components are divided by their values before the press perturbation and logged in order to bring all components to the same vertical scale. Shaded areas denote the total deviations to the stationary expectations. (C) Integrals across time of each community variability component. For all panels, **K**_**0**_ = (600, 400) ; *r*_1_ = *r*_2_ = 0.01 ; ***δ*** = (0.25, −0.25) ; *σ*_*e*_ = 0.01 and *α* = 0.1. Stat. Avg.: statistical averaging effect. Comp. Dyn.: compensatory dynamics effect.

This effect can be explained by the deviations in average population variability and statistical averaging effect (Figure 4B). Because average population variability is weighted by relative abundances, the deficit of population variability of the decreasing rarer species does not fully compensate for the excess of variability of the increasing and abundant species. Moreover, the evenness of species variances decreases faster than it would if it followed stationary expectations (Figure 4A, middle), yielding a deficit in statistical averaging effect (Zhao et al. 2022) which contributes positively to the deviation in community variability. The remaining component of community variability, the compensatory dynamics effect, is the component that reacts the less to the press perturbation, with a low negative value.

The interactions among the spread of the press perturbation within the community, the underlying community’s abundance distributions and the growth rate distributions generate rich patterns, even for simple two-species communities. Scenarii where the two species share the same response to the press perturbation generate the strongest absolute deviations in community variability, for “increase” and for “decrease” (Figure 5). In these settings, positive responses expectedly generate positive community variability deviations and negative responses generate negative community variability deviations. Since variance evenness does not change much during the transient dynamics, these deviations are mostly driven by deviations in average population variability. When the two species do not have the same growth rate, the population trend of the faster-growing, more abundant species happens slightly more quickly than that of the slower-growing, rarer species. This mismatch of the relative speeds of the species’ transients generate subtle changes in the evenness of species mean abundances and slightly stronger changes in the evenness of species variances, causing small deviations in statistical averaging effect (Figures S1 and S2).

**Figure 5:**
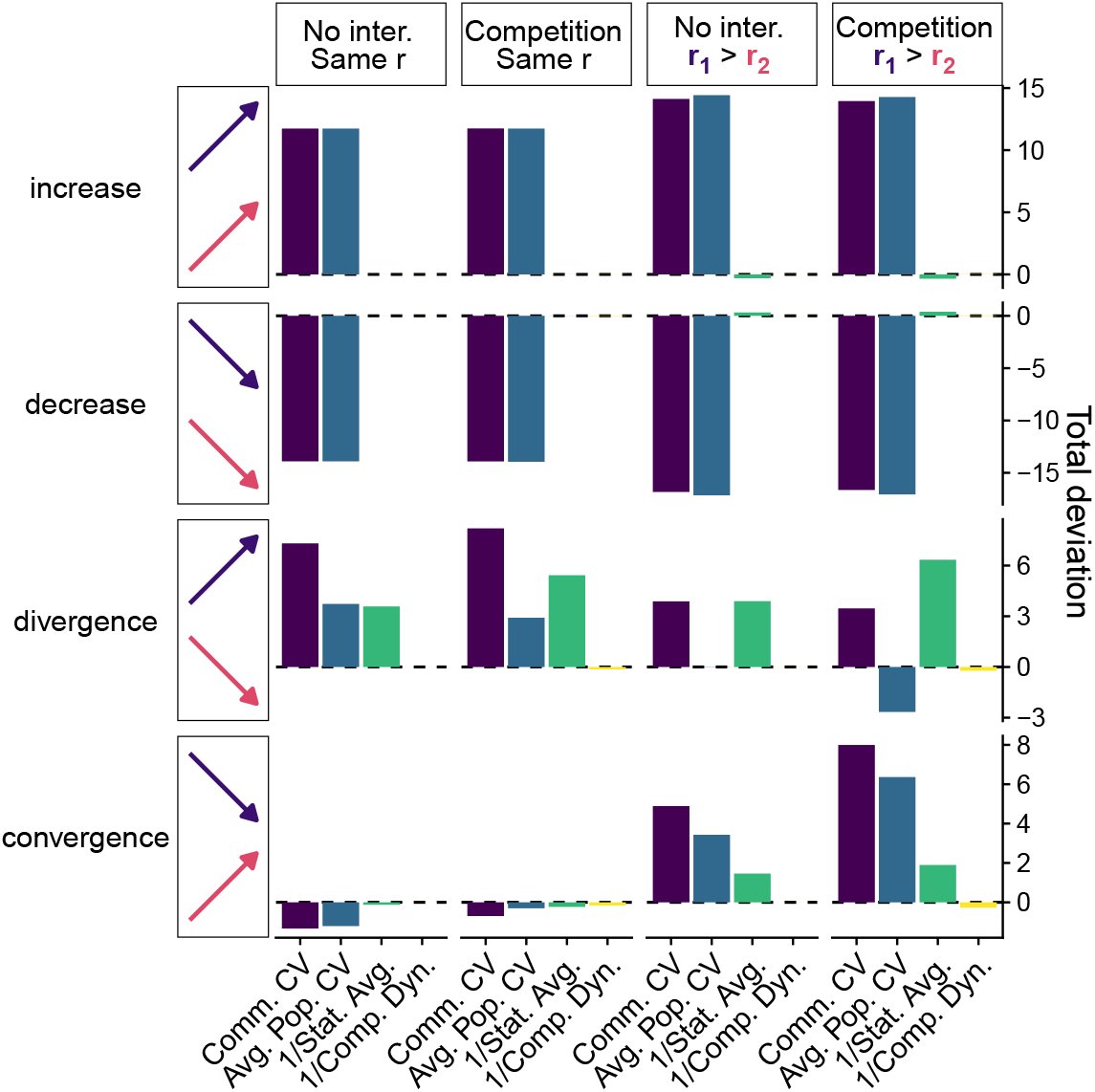
Impact of press perturbations (rows) on community variability during transient dynamics for different community contexts (columns). In each panel, total deviations from the stationary regime are reported for community variability and its components. Press perturbation scenarii are noted by colored arrows and represent different values of the relative perturbation vector ***δ***, namely from to top bottom: (0.25, 0.25), (−0.25, −0.25), (0.25, −0.25) and (−0.25, 0.25). For community contexts, **K**_**0**_ = (600, 400) for all panels, with varying interspecific competition strength (No inter.: *α* = 0; Competition: *α* = 0.1) and varying relative species growth rates (Same r: *r*_1_ = *r*_2_ = 0.01 ; r1 > r2: **r** = (0.01, 0.0066)). Stat. Avg.: statistical averaging effect. Comp. Dyn.: compensatory dynamics effect.

In the “convergence” and “divergence” scenarii, where responses to press are opposite between the two species, the deviations of community variability from the stationary expectation are weaker than in the other scenarii, regardless of their sign (Figure 5). In contexts where the two species share the same growth rate (left half of Figure 5), the community deviation is driven by the population trend of the most abundant species, via both the deviation in average population variability and the deviation in statistical averaging effect. In the “convergence” scenario, deviations in community variability components are negative but small in magnitude, because the deviations in average population variability and in statistical averaging effect change sign across time (Figure S3, top half). This is due to the community crossing the point at which the rarer species, because of the press perturbation, becomes the more abundant one.

Adding interspecific competition when the two species have the same growth rates, the deviation in average population variability is decreased in the “divergence” scenario while the deviation in inverse statistical averaging is increased (left half of Figure 5). Conversely, in the “convergence” scenario, adding interspecific competition increases the deviation of average population variability while the deviation in inverse statistical averaging stays similar. Inspecting the temporal change of each variability component (Figs. S3-S4) reveals that this arises because, with interspecific competition, in the later part of the transient dynamics the components take more time to reach their stationary values than without competition. When a component’s deviation keeps the same sign across the transient regime, its deviation is thus amplified (*e*.*g*. statistical averaging in Fig. S3). Conversely, when a component’s deviation in the beginning of the transient is opposite to its deviation in the end of the transient regime, the deviation is amplified (resp. buffered) by interactions if the end (resp. the beginning) of the transient regime dominates the integral. See for example the contrast between inverse statistical averaging in the first and in the second columns of Figure S3 (amplification) as opposed to the contrast between average population variability in the first and in the second columns of Figure S4 (buffering). Small deviations in compensatory dynamics effect occur only in the presence of interspecific competition, and always contribute negatively to the deviation in community variability.

In the community contexts with different growth rates across species (right half of Figure 5), the rarer, slower-growing species takes more time to reach its stationary mean than the more abundant, faster-growing species. This increases the contribution of the rarer species to the average population variability deviation, increasing total deviation in community variability in the “turnover” scenario but reducing it in the “divergence” scenario. Again, interspecific competition exacerbates the patterns in the end of the transient regimes. Contrastingly to all other cases, in the right half of the third row of Figure 5, total deviations in average population variability and in inverse statistical averaging effect are both noticeable but have opposed signs. There, the positive deviation in inverse statistical averaging explains the positive deviation in community variability (see Figure S3).

## Discussion

Our results reveal that the temporal variability of populations and communities subjected to a press perturbation as well as short-term environmental stochasticity deviates during transient dynamics from what could have been expected near equilibrium, under environmental stochasticity alone. Moreover, the distribution of the responses to the press perturbation among species within communities alters how community variability deviates from its stationary expectation, depending on which of the abundant or rare species is the one that increases or decreases. While these results are well in line with the framework that variability reflects a system’s inherent stability in a given perturbation context (Arnoldi et al. 2019), our results clearly illustrate how the response to press perturbations interacts with the response to environmental stochasticity. Especially, the dependence of population variability deviation to the direction of the long-term trend was not predicted by classical methods that linearize community dynamics around a fixed point. Since we show that long-term changes in mean abundance impact the variability around these changes, these two dimensions of ecological stability appear fundamentally related (Donohue et al. 2013).

Our findings at the scale of population variability resonate with increasing efforts to consider transient dynamics in population biology (Ezard et al. 2010). By estimating population projection matrix models for many wild species and comparing the corresponding stability metrics with the species’ long-term growth, Gamelon et al. (2014) found that declining populations may be more efficient at buffering disturbances than increasing populations. This corroborates our findings in a different conceptual and methodological framework (Stott et al. 2011). An intuition for explaining this phenomenon lies in the asymmetry of the stability landscape of the logistic growth around the carrying capacity (Dennis et al. 2016). Above carrying capacity (and thus during a decline), the stability landscape is steeper, pulling the population to its equilibrium faster than if it was below carrying capacity. This was never detected in studies using the near-equilibrium assumption because linearization around the carrying capacity implies a symmetric stability landscape.

We predict that in communities where species share a common negative response to a press perturbation, such as microcosm warming experiments (Pennekamp et al. 2018), measured community variability could reflect lower values than expected near equilibrium. Our results show that such situations pose a risk of overestimating ecological stability for such communities, or underestimating the strength of environmental stochasticity.

Extending our approach to more complex communities is key in understanding when strong deviations in population and community variability can be expected. Just like the number of species generally dampens community variability via an increase in the diversity of responses to environmental fluctuations (Loreau and de Mazancourt 2013; Danet et al. 2025; Polazzo et al. 2025), we expect that increasing the number of species will dampen the deviation of community variability to its stationary expectation via an increase in the diversity of responses to the press perturbation. Even in that case, quantifying deviations from stationary expectations would remain helpful to investigate the relationship between the number of species and the average population variability (Xu et al. 2021; Danet et al. 2025). Interspecific competition has been shown to affect community variability (Loreau and de Mazancourt 2008) and asymptotic response to press perturbations (Lajaaiti et al. 2025). Here we illustrate that competition also affects transient dynamics by slowing down the long-term trends in the end of the response to the press perturbation, altering community variability deviation. Extending our approach to different types of interactions and community structures could help uncover how long-term changes cascading through ecological networks could affect variability measurements (Duchenne et al. 2020; Danet et al. 2021).

Species long-term trends are associated to specific functional trait combinations that determine species responses to anthropogenic pressures (Henn et al. 2024). Notably, from metabolic scaling and evolutionary trade-offs, species that reproduce faster tend to exhibit higher population abundances while also having smaller body size (Brown et al. 2004). Moreover, large body size is linked to high extinction risk (Chichorro et al. 2019). The population abundance of large-bodied species is thus expected (i) to be lower on average, (ii) to be more variable because they have a lower growth rate and (iii) to be more likely to decline. Many natural communities are thus expected to follow the “divergence” scenario of our study, where community variability is always higher than expected near equilibrium (Figure 5). For some taxa, other traits than body size may explain the response to pressures, such as insectivory in the case of wild birds (Bowler et al. 2019) or habitat specialization generally (Chichorro et al. 2019). Identifying the traits responsible for ecological stability is at the core of bridging advances in functional ecology with ecological stability theory (Lavorel and Garnier 2002; de Bello et al. 2021). By showing that deviations in community variability can arise from the co-occurence of traits associated to long-term trends and traits associated to variability, we support this framework across perturbation types.

Whether or not these deviations in population and community variability could actually be detected in biodiversity survey data and correctly attributed to long-term population changes is not easily predictable. Firstly, our predictions only consider press perturbations as abrupt changes in carrying capacity, while real-world perturbations applied to ecological systems also change in time (Díaz and Malhi 2022; Jaureguiberry et al. 2022), and may affect not only to the carrying capacity but also the strength of environmental stochasticity itself (Seddon et al. 2016; Vázquez et al. 2017). Secondly, even for population variability, the deviations from stationary expectations that we observe have a small magnitude. Since measuring variability out of equilibrium is a challenge in itself (Clark et al. 2022), detecting changes in variability may be even more difficult, especially for systems under different stochasticity types (Engen et al. 1998; Arnoldi et al. 2019). In supporting analyses, we extend our methods to demographic and migration stochasticity, and we show that our results for environmental stochasticity also hold for demographic stochasticity (supplemental Section S3, Fig. S5).

Moment closure approximations in ecology have first been introduced in birth-death models, where the probability of demographic events is modelled explicitly (Nåsell 2003; Newman et al. 2004). They prove a valuable tool for studying stochastic systems, even if they remain approximations (see Marrec et al. 2023), which could be made more accurate by also modelling the temporal dynamics of the skewness rather than only the mean and variance (Nåsell 2003). In our methods, the moment closure approximation holds only when stochasticity strength and growth rate are not too high, which is an expected limitation. Simulation studies could help better quantifying the magnitude of the error associated to our approximation, but may hit the limitation that numerical methods for integrating stochastic differential equations also rely on approximations (e.g. Rößler 2010). Aside from generating useful insights for ecological variability, moment closure approximations also generate predictions for the means of ecological properties and could help further reveal how stochasticity shapes ecological systems, near or far from equilibrium (Shoemaker et al. 2020).

The importance of transient phenomena in ecology has been proposed for a long time (Hastings 2004), but their study has been restricted to a few particular settings such as systems with alternative stable states (Scheffer 2009; Hastings et al. 2018). We show that out-of-equilibrium dynamics can also matter for systems usually studied near equilibrium, and where transient dynamics are generated by abrupt changes in parameter values.

Overall, our results reveal a mathematical relationship between two dimensions of ecological stability and call for further investigation of the impact of long-term population trends on population and community variability. Under multiple perturbation regimes at multiple time scales (Hastings 2016), the fact that community variability integrates intrinsic ecological properties with the strength of several perturbation types is an avenue towards building biodiversity indicators that accurately reflect the intensity of anthropogenic perturbations across ecological and geographical scales.

## Acknowledgements

We thank Jean-François Arnoldi for valuable advice on mathematical methods. This work was supported by Agence Nationale de la Recherche (ANR) under grant number ANR-23-CE02-0030.

## Author Contribution Statement

JG, DG, CF and ET conceptualized the study. DG and JG conducted mathematical analyses. DG wrote Python code and JG wrote Julia code. DG and JG visualized the results. JG wrote the original draft and all authors contributed to review and editing.

## Code Accessibility Statement

All code required to reproduce the results as well as simulated data are available on Zenodo (access link available on request to the corresponding author) and in a GitHub repository (will be made public after acceptance).

