## Supplemental Information for "Population and community variability deviate from stationary expectations during transient dynamics"

#### S1 Mean and variance of stochastic differential equations

In this appendix section, we detail how stochastic calculus theory is used to build deterministic approximation for the statistical moments of the solution of stochastic differential equations.

##### *S1.1 Previous mathematical results*

**Definitions.** Let $\mathbf{X_t}$ a stochatic (Itô) process of dimension $S$. Let $p_X(\mathbf{X}, t)$ the probability density of $\mathbf{X_t}$ at time $t \in \mathbb{R}^+$. Let $X_{t,i}$ the $i$-th component of $\mathbf{X_t}$.

Let $\boldsymbol{\mu}_t = \mathbb{E}[\mathbf{X_t}]$ the expectation of $\mathbf{X_t}$ and let $\boldsymbol{\Sigma}_t = \mathbb{E}[(\mathbf{X_t} - \boldsymbol{\mu}_t)(\mathbf{X_t} - \boldsymbol{\mu}_t)^T]$ the $S$ by $S$ variance-covariance matrix of $\mathbf{X_t}$.

$\nabla$ is the vector differential operator, *i.e.* for $\phi$ a (differentiable) function of $S$ variables $(x_1, \dots, x_S)$, $\nabla\phi = (\frac{\partial\phi}{\partial x_1}, \dots, \frac{\partial\phi}{\partial x_S})$.

We consider the stochastic differential equation (SDE) in Eq. 1:

$$\mathrm{d}\mathbf{X_t} = f(\mathbf{X_t}, t)\mathrm{d}t + G(\mathbf{X_t}, t)\mathrm{d}\boldsymbol{W_t} \tag{S1}$$

$f$ is a twice differentiable vector-valued function, $G$ is a matrix-valued function. $p_X(\mathbf{X}, t)$ satisfies the Fokker-Planck-Kolmogorov equation, which is a partial differential equation. While it is exact, it is not easily solvable even for simple functions $f$ and $G$ (Särkkä and Solin 2019). In the following, we present existing results that allow to build an approximation for the change in time of the statistical moments of $\mathbf{X_t}$.

**Itô's lemma.** Let $\phi(\mathbf{X_t}, t)$ any scalar function of the process $\mathbf{X_t}$. The Itô stochastic differential equation for $\phi$ is given by:

$$\mathrm{d}\phi = \frac{\partial \phi}{\partial t}\mathrm{d}t + (\nabla\phi)^T\mathrm{d}\mathbf{X_t} + \frac{1}{2}\,\mathrm{tr}[(\nabla\nabla^T\phi)\mathrm{d}\mathbf{X_t}\mathrm{d}\mathbf{X_t}^T] \tag{S2}$$

This equation can then be expanded using Equation S1 with the following rules for mixed differentials: $\mathrm{d}\mathbf{W_t}\ \mathrm{d}t = \mathbf{0}$ ; $\mathrm{d}t\ \mathrm{d}\mathbf{W_t} = \mathbf{0}$ ; $\mathrm{d}\mathbf{W_t}\mathrm{d}\mathbf{W_t}^T = \mathrm{d}t$.

**General formula.** Applying Itô's lemma on Equation S1 with $\phi(\mathbf{X_t}) = X_{t,i}$ (for elements of $\boldsymbol{\mu_t}$) and $\phi(\mathbf{X_t}) = X_{t,i}X_{t,j} - \mu_{t,i}\mu_{t,j}$ (for elements of $\boldsymbol{\Sigma_t}$) and taking the expectation yields (Särkkä and Solin 2019):

$$\begin{aligned}\frac{\mathrm{d}\boldsymbol{\mu_t}}{\mathrm{d}t} &= \mathbb{E}[f(\mathbf{X_t}, t)] \\ \frac{\mathrm{d}\boldsymbol{\Sigma_t}}{\mathrm{d}t} &= \mathbb{E}[f(\mathbf{X_t}, t)(\mathbf{X_t} - \boldsymbol{\mu_t})^T] + \mathbb{E}[(\mathbf{X_t} - \boldsymbol{\mu_t})f(\mathbf{X_t}, t)^T] + \mathbb{E}[G(\mathbf{X_t}, t)G(\mathbf{X_t}, t)^T]\end{aligned} \tag{S3}$$

When $f$ is non-linear, the first two terms of the time-derivative of $\mathbf{\Sigma}_t$ can depend not only on $\mathbb{E}[\mathbf{X_t}]$ and $\mathbb{E}[\mathbf{X_t}^2]$, but also on statistical moments of $\mathbf{X_t}$ of higher order than 2, which time-derivatives then depend on even higher order moments, and so on.

**Moment closure approximation.** To close this relationship, we use a first-order linearization of $f$ around $\boldsymbol{\mu}_t$ in the expression for the time-derivative of $\mathbf{\Sigma}_t$ (Särkkä and Solin 2019). $f(\mathbf{X_t}, t)$ is approximated by $f(\boldsymbol{\mu}_t, t) + \mathbf{F}(\boldsymbol{\mu}_t, t)(\mathbf{X_t} - \boldsymbol{\mu}_t)$ if $\mathbf{F}(\boldsymbol{\mu}_t, t)$ is the Jacobian matrix of $f(., t)$ evaluated in $\boldsymbol{\mu}_t$, and Equation S3 thus yields:

$$\begin{aligned} \frac{\mathrm{d}\boldsymbol{\mu}_t}{\mathrm{d}t} &= \mathbb{E}[f(\mathbf{X_t}, t)] \\ \frac{\mathrm{d}\mathbf{\Sigma}_t}{\mathrm{d}t} &\approx \mathbb{E}[\mathbf{F}(\boldsymbol{\mu}_t, t)(\mathbf{X_t} - \boldsymbol{\mu}_t)(\mathbf{X_t} - \boldsymbol{\mu}_t)^T] + \mathbb{E}[(\mathbf{X_t} - \boldsymbol{\mu}_t)(\mathbf{F}(\boldsymbol{\mu}_t, t)(\mathbf{X_t} - \boldsymbol{\mu}_t))^T] + \\ &\quad \mathbb{E}[G(\mathbf{X_t}, t)G(\mathbf{X_t}, t)^T] \\ &= \mathbf{F}(\boldsymbol{\mu}_t, t)\mathbf{\Sigma}_t + \mathbf{\Sigma}_t\mathbf{F}(\boldsymbol{\mu}_t, t)^T + \mathbb{E}[G(\mathbf{X_t}, t)G(\mathbf{X_t}, t)^T] \end{aligned} \tag{S4}$$

Note that this first-order approximation relies on the assumption that $\mathbf{X_t}$ stays relatively close to $\boldsymbol{\mu}_t$. In our study cases, since $G$ never includes powers of $\mathbf{X_t}$ higher than 1, expressions of $\mathbb{E}[G(\mathbf{X_t}, t)G(\mathbf{X_t}, t)^T]$ never require high order moments of $\mathbf{X_t}$ and therefore we can keep their exact expression even in the approximated system.

**Stationary regime.** $\mathbf{X_t}$ reaches a stationary regime when its probability distribution is constant in time. In the moment closure approximation, the fixed points $(\boldsymbol{\mu}^*, \mathbf{\Sigma}^*)$ of system S4, *i.e.* the values for means and variance-covariance matrix elements for which $\frac{\mathrm{d}\boldsymbol{\mu}_t}{\mathrm{d}t} = \frac{\mathrm{d}\mathbf{\Sigma}_t}{\mathrm{d}t} = 0$, are approximations to the statistical moments of the stationary distribution of $\mathbf{X_t}$ (Särkkä and Solin 2019).

#### *S1.2 Applications to ecological modelling*

**Stochastic logistic growth.** $X_t$ is a scalar representing the population abundance of a single species, $f(X_t, t) = f(X_t) = rX_t(1 - \frac{X_t}{K})$, and $\boldsymbol{\mu_t}$, $\boldsymbol{\Sigma_t}$ and $G(X_t, t)$ are scalars written $\mu_t$, $V_t$ and $g(X_t)$, respectively. Under environmental stochasticity, $g(X_t) = \sigma_e X_t$ (Arnoldi et al. 2019) and $\mu_t$ and $V_t$ thus satisfy equation 3. See Section S3 for different types of stochasticity.

**Generalized Lotka-Volterra system.** With $f$ set according to the generalized Lotka-Volterra model with interaction matrix $\mathbf{A}$ (Equation 5) and under environmental stochasticity for $G$, Equation S4 becomes:

$$\begin{aligned}\frac{\mathrm{d}\boldsymbol{\mu_t}}{\mathrm{d}t} &= \mathrm{diag}(\boldsymbol{\mu_t})(\mathbf{r} + \mathbf{A}\boldsymbol{\mu_t}) + \mathrm{diag}(\boldsymbol{A\Sigma_t}) \\ \frac{\mathrm{d}\boldsymbol{\Sigma_t}}{\mathrm{d}t} &= \mathbf{F}(\boldsymbol{\mu_t})\boldsymbol{\Sigma_t} + \boldsymbol{\Sigma_t}\mathbf{F}(\boldsymbol{\mu_t})^T + \sigma_e^2\,\mathrm{diag}[\mathrm{diag}(\boldsymbol{\mu_t})\boldsymbol{\mu_t} + \mathrm{diag}(\boldsymbol{\Sigma_t})]\end{aligned} \tag{S5}$$

Here, when $\mathbf{v}$ is a vector, we note $\mathrm{diag}(\mathbf{v})$ the matrix whose diagonal elements are the elements of $\mathbf{v}$ and when $\mathbf{M}$ is a matrix, $\mathrm{diag}(\mathbf{M})$ is the vector whose elements are the elements of the principal diagonal of $\mathbf{M}$. Thus $\mathrm{diag}(\mathrm{diag}(\mathbf{v})) = \mathbf{v}$ but $\mathrm{diag}(\mathrm{diag}(\mathbf{M}))$ replaces all off-diagonal terms of $\mathbf{M}$ by zero.

$\mathbf{F}(\boldsymbol{\mu_t})$ is the Jacobian matrix of $f$ evaluated at $\boldsymbol{\mu_t}$. Setting all diagonal elements of $\mathbf{A}$ to $-1$ (see Methods), $F_{ii}(\boldsymbol{\mu}) = r_i(1 - 2\frac{\mu_i}{K_i})$ and $F_{ij}(\boldsymbol{\mu}) = A_{ij} r_i \frac{\mu_i}{K_i}$.

From numerical solutions of Equation S5, mean community abundance $\mu_c$ and expected community variance $V_c$ can each be computed for any time point as the sum of all elements of $\mu_\mathbf{t}$ and of $\boldsymbol{\Sigma_t}$, respectively.

### S2 Supplemental analyses

#### S2.1 Analytical results for the stochastic logistic growth

**Stationary regime.** Setting $\frac{\mathrm{d}\mu_t}{\mathrm{d}t} = \frac{\mathrm{d}V_t}{\mathrm{d}t} = 0$ in equation 3 and solving for $(\mu^*, V^*)$ yields three possible fixed points. The first is the trivial fixed point $(\mu^*, V^*) = (0, 0)$.

If $\mu^* \neq 0$ and $V^* \neq 0$, we get:

$$\begin{cases} V^* = \mu^*(K - \mu^*) \\ 0 = 2\mu^{*2} - 3K\mu^* + K^2(1 + \dfrac{\sigma_e^2}{2r}) \end{cases} \tag{S6}$$

Therefore $\mu^*$ has two candidate values, $\mu^* = \frac{3}{4}K \pm \frac{1}{4}K\sqrt{1 - 8\frac{\sigma_e^2}{2r}}$. Nåsell (2003) show that the smallest of these values can be ruled out as spurious (i.e. it is not a possible stationary mean for the stochastic process), which leaves the fixed point where $\mu^*$ is close to but still smaller than $K$. When $\sigma_e^2$ is too high, the system approaches a saddle-node bifurcation where the fixed points vanish.

**Isoclines equations.** From above, the isocline for the mean satisfies $\frac{\mathrm{d}\mu_t}{\mathrm{d}t} = 0 \iff V = \mu(K - \mu)$ (red thick line on Fig. 2B).

For the variance isocline (blue thick line on Fig. 2B),

$$\begin{aligned} \frac{\mathrm{d}V_t}{\mathrm{d}t} = 0 &\iff V(2r(2\frac{\mu}{K} - 1) - \sigma_e^2) = \sigma_e^2\mu^2 \\ &\iff V = \mu^2\frac{\sigma_e^2}{2r(2\frac{\mu}{K} - 1) - \sigma_e^2} \end{aligned} \tag{S7}$$

If $\mu = \mu^*$, this formulation for $V^*$ makes it easier to see that $V^* \approx \mu^{*2}\frac{\sigma_e^2}{2r-\sigma_e^2}$ than by substituting the analytical expression for $\mu^*$ in Equation S6.

#### S2.2 *Comparison between the stationary regime and the isocline for variance*

Here we show that the relative positions of the stationary expectation for variance (dashed line) and of the variance isocline (blue line) always matches Figure 2.

We consider a trajectory $\{(\mu_t, V_t)\}_{t>t_d}$ where the mean and variance of the stochastic logistic growth converge towards their stationary regime $(\mu^*, V^*)$.

For any mean value $\mu_t$, the stationary expectation for variance, $\hat{V}(\mu_t)$, is the variance that the system would have if it was in a stationary regime at mean $\mu_t$. It is obtained by setting $\frac{\mathrm{d}V_t}{\mathrm{d}t} = \frac{\mathrm{d}\mu_t}{\mathrm{d}t} = 0$ in Equation 3 and solving it for $(\hat{K}(\mu_t), \hat{V}(\mu_t))$, with $\hat{K}(\mu_t)$ the carrying capacity required to observe a stationary regime at mean $\mu_t$.

Using the result of section S2.1, we can also define $V_{iso}(\mu_t)$, the nullcline for variance, *i.e.* the value of $V_t$ for which $\frac{\mathrm{d}V_t}{\mathrm{d}t} = 0$. Graphical analysis shows that $\frac{\mathrm{d}V_t}{\mathrm{d}t} > 0$ when $V_t > V_{iso}(\mu_t)$. Notice that, by definition, $\hat{V}(\mu^*) = V_{iso}(\mu^*)$.

We need to prove that, when the mean is above (resp. below) its stationary value, the variance isocline is below (resp. above) the stationary expectation. We detail the proof for a negative long-term trend, *i.e* we show that $\mu_t > \mu^* \implies V_{iso}(\mu_t) < \hat{V}(\mu_t)$.

Let $\alpha$ such that $\mu^* = K\alpha$. According to the solution of Equation S6, $\alpha = 3/4 + 1/4\sqrt{1 - \frac{4\sigma^2}{r}}$ and thus $3/4 < \alpha < 1$. Also note that according to the same system, $\mu_t = \hat{K}(\mu_t)\alpha$.

Thus $\mu_t > \mu^* \implies \hat{K}(\mu_t)\alpha > K\alpha \implies \hat{K}(\mu_t) > K$.

From Equation S7, $\hat{V}(\mu_t)(2r\ (2\frac{\mu_t}{\hat{K}(\mu_t)} - 1) - \sigma_e^2) = \sigma_e^2\mu_t^2 = V_{iso}(\mu_t)(2r(2\frac{\mu_t}{K} - 1) - \sigma_e^2)$. If $\hat{K}(\mu_t) > K$ then, since all coefficients are strictly positive, $V_{iso}(\mu_t) < \hat{V}(\mu_t)$.

#### S2.3 Supplemental figures

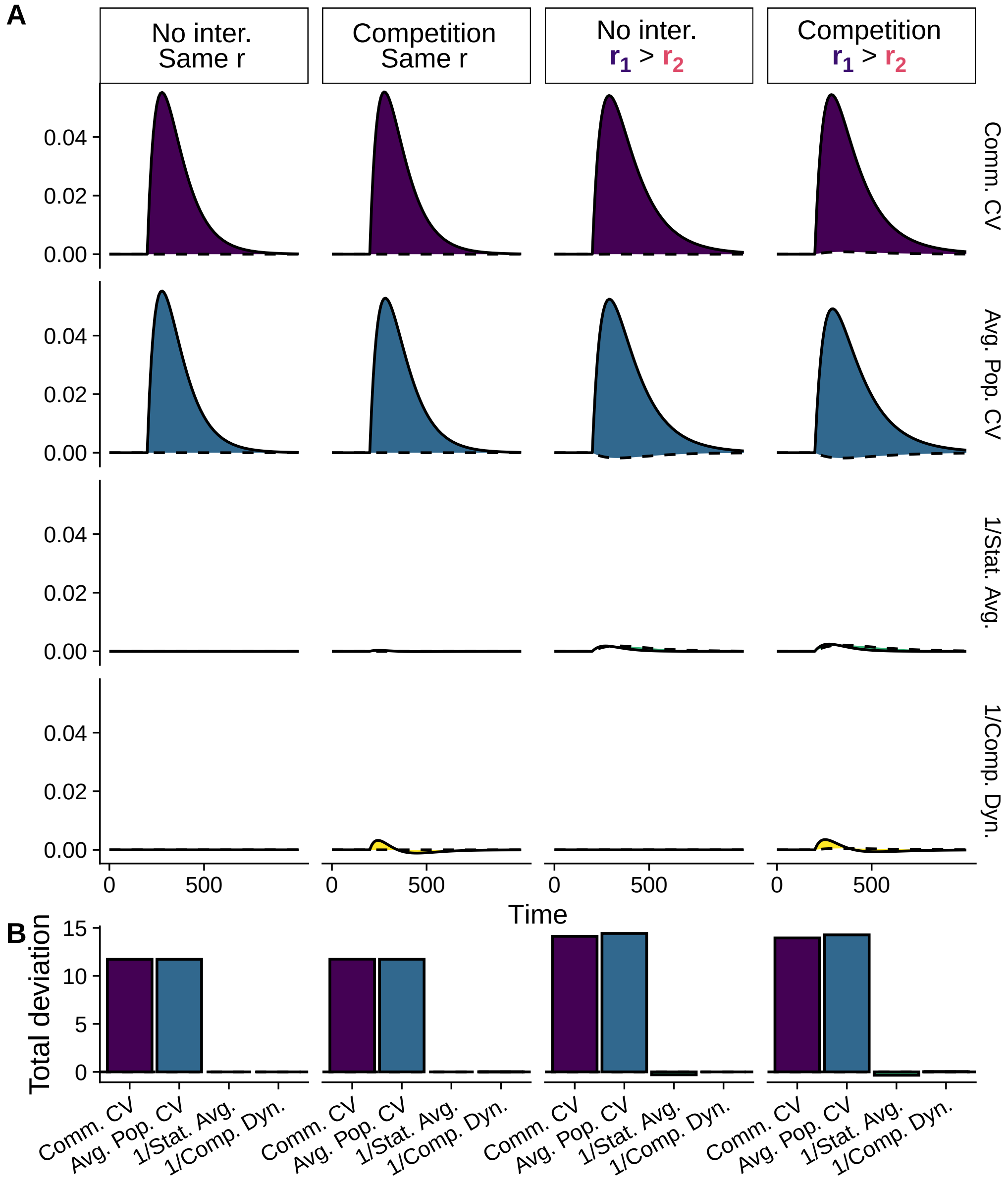


Figure S1: **Detail on the impact of the "increase" press perturbation in different community contexts.** Time series and total deviations correspond to the first row of Figure 5. In the "increase" scenario, both species increase under the press perturbation, *i.e.* $\delta = (0.25, 0.25)$. Community contexts (different columns) correspond to the different columns in Figure 5.

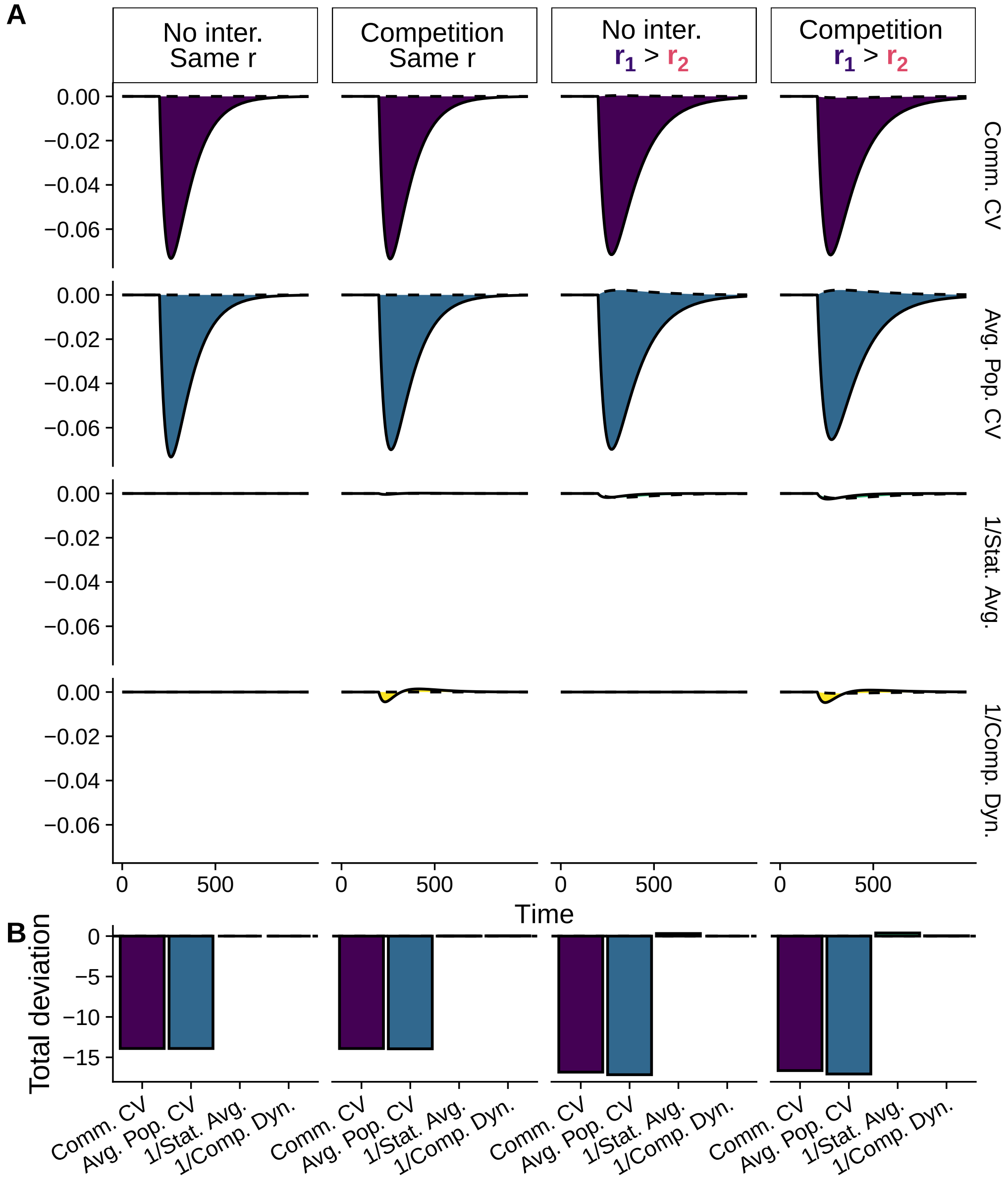


Figure S2: **Detail on the impact of the "decrease" press perturbation in different community contexts.** Time series and total deviations correspond to the second row of Figure 5. In the "decrease" scenario, both species decline under the press perturbation, *i.e.* $\delta = (-0.25, -0.25)$. Community contexts (different columns) correspond to the different columns in Figure 5.

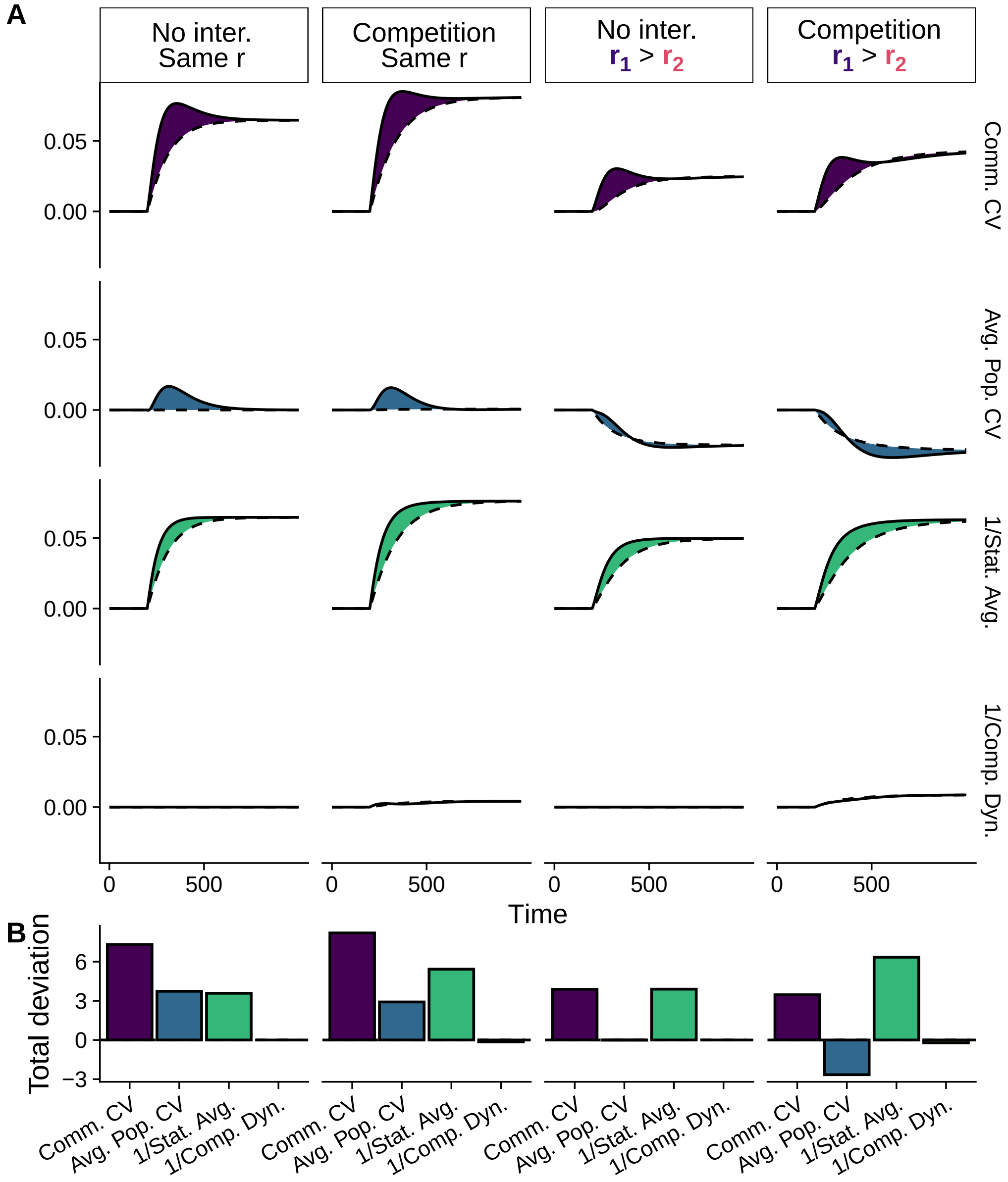


Figure S3: **Detail on the impact of the "divergence" press perturbation in different community contexts.** Time series and total deviations correspond to the third row of Figure 5. In the "divergence" scenario, the most abundant species increases and the rarer species declines, *i.e.* $\delta = (0.25, -0.25)$. Community contexts (different columns) correspond to the different columns in Figure 5.

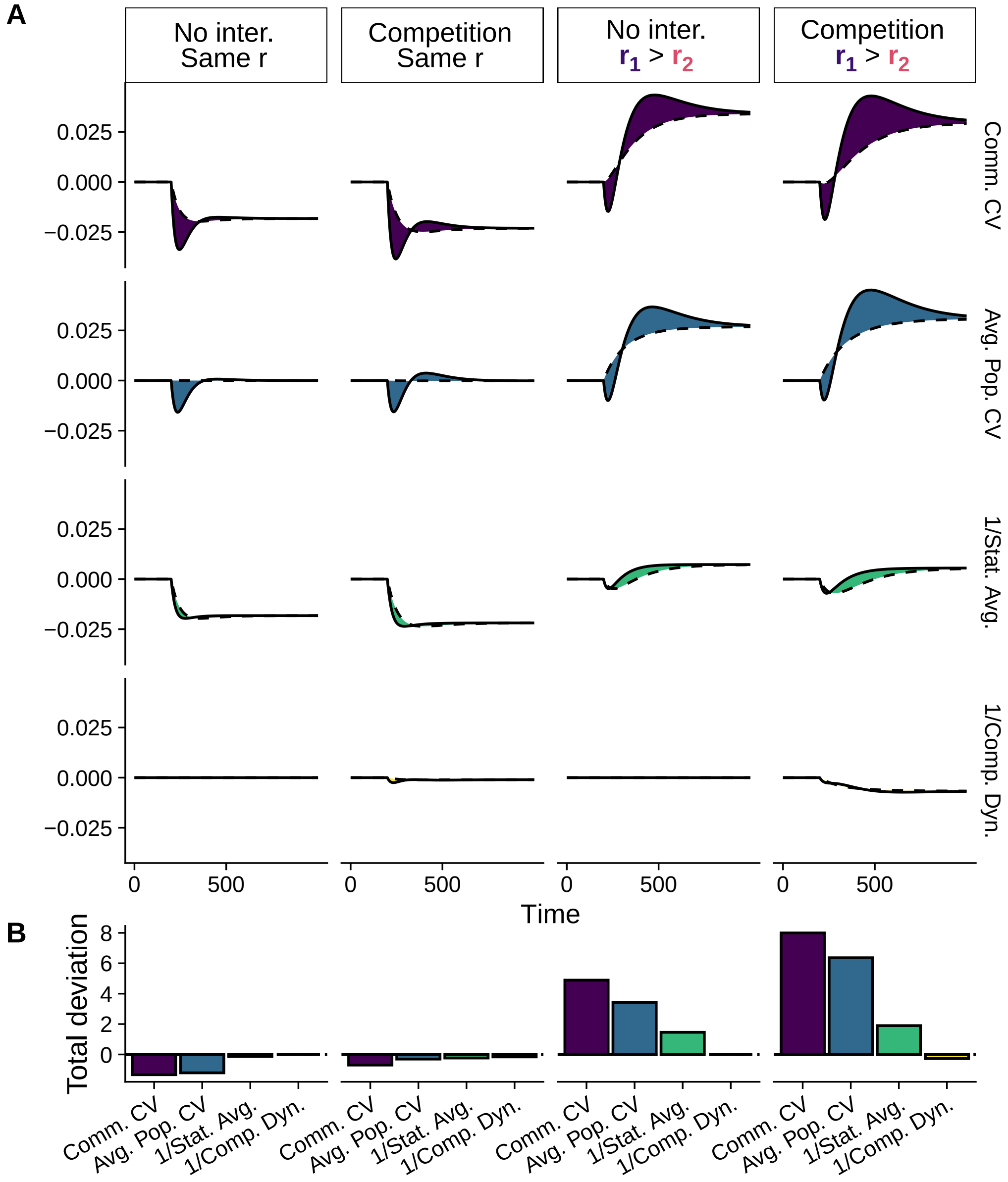


Figure S4: **Detail on the impact of the "convergence" press perturbation in different community contexts.** Time series and total deviations correspond to the last row of Figure 5. In the "convergence" scenario, the most abundant species declines and the rarer species increases, *i.e.* $\delta = (-0.25, 0.25)$. Community contexts (different columns) correspond to the different columns in Figure 5.

#### S3 Extensions to different stochasticity types

Following Arnoldi et al. (2019), Equation 1 can translate several types of stochasticity by introducing a parameter $\gamma$, the stochasticity type, in order to change the scaling of the diffusion function with population abundance. Namely, for independent species-level perturbations, $G_\gamma(\mathbf{X_t}, t)_i = \sigma_\gamma X_i(t)^\gamma$. For environmental stochasticity, $\gamma = 1$ (see Methods). Setting $\gamma = \frac{1}{2}$ corresponds to demographic stochasticity and $\gamma = 0$ to stochasticity due to immigration and emigration (Arnoldi et al. 2019).

##### *S3.1 Relationship between stationary mean and variance*

For a single population, Equation S4 yields:

$$\begin{aligned}\frac{\mathrm{d}\mu_t}{\mathrm{d}t} &= r\mu_t\left(1 - \frac{\mu_t}{K}\right) - \frac{r}{K}V_t \\ \frac{\mathrm{d}V_t}{\mathrm{d}t} &= 2rV_t(1 - 2\frac{\mu_t}{K}) + \sigma_\gamma^2\mathbb{E}[X_t^{2\gamma}]\end{aligned} \tag{S8}$$

For $\gamma = 1/2$, and solving Equation S8 for the stationary regime yields $\mathbb{E}[X_t^{2\gamma}] = \mu_t$ and $V^* \approx \mu^* \frac{\sigma_d^2}{2r}$. For $\gamma = 0$, $\mathbb{E}[X_t^{2\gamma}] = 1$ and $V^* \approx \frac{\sigma_m^2}{2r}$.

Standardizing population variance by population mean thus must take a different shape depending on the type of stochasticity, for which we propose a coefficient of variation that depends on $\gamma$:

$$\mathrm{CV}_\gamma = \frac{\sqrt{V}}{\mu^\gamma} \tag{S9}$$

Note that, under environmental stochasticity, $\text{CV}_\gamma$ is the classical coefficient of variation used in the main manuscript.

#### S3.2 Decomposing the generalized coefficient of variation

In the framework of Thibaut and Connolly (2013), the (classical) coefficient of variation of community abundance can be decomposed as the product of the average population coefficient of variation with the population synchrony index of Loreau and de Mazancourt (2008), $\phi$. In the following, we derive the same decomposition for any positive value of $\gamma$.

Now with $V_c$ the variance of the abundance a community composed of $S$ species under stochasticity type $\gamma$, with $\mu_c = \sum_{i=1}^{S} \mu_i$ the average of community abundance. $V_i$ denotes the variance of the abundance of species $i$. Loreau and de Mazancourt (2008) define the species synchrony $\phi$ as:

$$V_c = \phi(\sum_{i=1}^{S} \sqrt{V_i})^2 \tag{S10}$$

Taking the square root of Equation S10 and dividing by $\mu_C^\gamma$, the generalized coefficient of variation of the community's abundance is:

$$\text{CV}_{\gamma,c} = \sqrt{\phi} \sum_i \frac{\sqrt{V_i}}{\mu_C^\gamma}$$

Following Thibaut and Connolly (2013), multiplying the fraction inside the sum by $\mu_i^\gamma/\mu_i^\gamma$ yields:

$$\text{CV}_{\gamma,c} = \sqrt{\phi} \sum_i \left(\frac{\mu_i}{\mu_c}\right)^\gamma \frac{\sqrt{V_i}}{\mu_i^\gamma}$$

Writing $p_i$ the relative (average) abundance of species $i$ in the community, i.e. $p_i = \frac{\mu_i}{\mu_c}$, we get:

$$\text{CV}_{\gamma,c} = \sqrt{\phi} \sum_i p_i^\gamma \text{CV}_{\gamma,i} \tag{S11}$$

The average population variability from Thibaut and Connolly (2013) thus has different weights depending on stochasticity type. Note that under environmental stochasticity, $\gamma = 1$ and Equation S11 is the result of Thibaut and Connolly (2013).

Statistical averaging effect and compensatory dynamics effects from Zhao et al. (2022) (see Methods) depend only on population and community variances, not on means, and can thus be used regardless of the stochasticity type.

#### *S3.3 Long-term trends under different stochasticity types*

Because of the varying importance of $p_i$ for the average population coefficient of variation, rare species increasingly drive the (generalized) community coefficient of variation as stochasticity type goes from environmental, to demographic, to migration. The population-level coefficients of variation of rare species are indeed decreasingly weighted by their low relative abundance, increasing their contribution.

In order to illustrate this, for the most simple community context of Figure 5, we vary the stochasticity type (and the formulae for community variability components) for each of the four press perturbation scenarii (Figure S5). For example, in the “divergence” scenario under migration stochasticity, the population variability deficit of the decreasing rare species completely compensates the population variability excess of the increasing abundant species. This is less the case for demographic stochasticity, and even less for environmental stochasticity (Figure S5). The increase and decrease scenarii are not affected by stochasticity type.

This result is already known near equilibrium (Arnoldi et al. 2019), but must be taken into consideration before applying our framework to real biodiversity data: the main stochasticity type can yield different dependences of community variability to the way long-term trends are spread between the distributions of species abundances and growth rates.

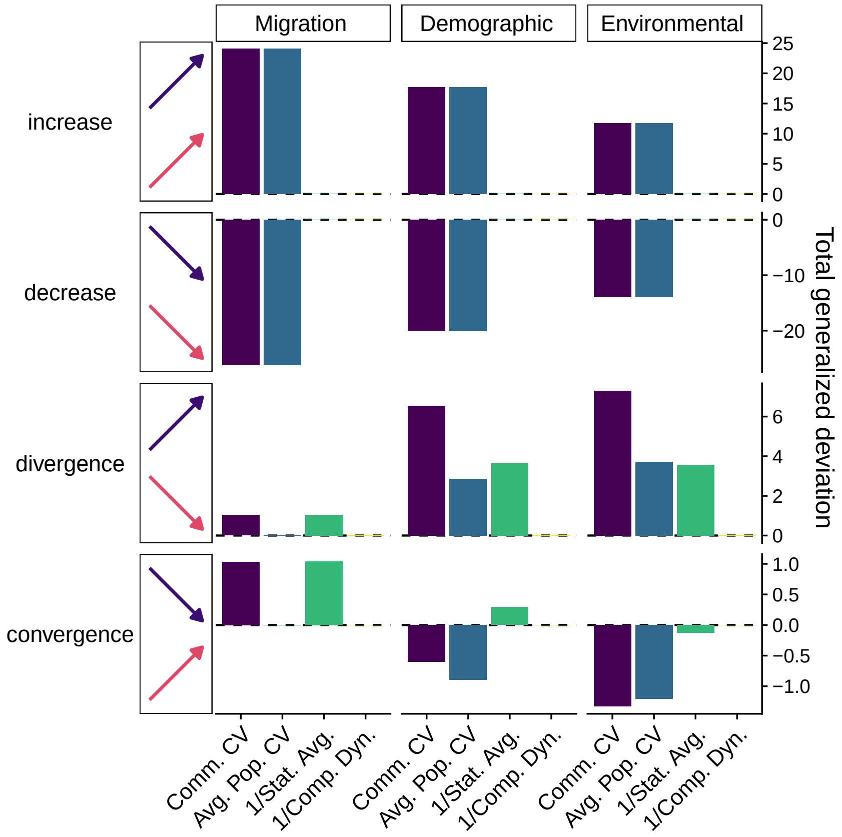


Figure S5: **Impact of stochasticity type on total deviation from stationary expectation.** The most simple community context is shown, with $\mathbf{K_0} = (600, 400)$; $r_1 = r_2 = 0.01$ and $\alpha = 0$. $\sigma_\gamma$ is set to 0.01 for all stochasticity types. Average population coefficient of variation is computed according to Equation S11.
